# Fog and live fuel moisture in coastal California shrublands

**DOI:** 10.1101/171850

**Authors:** Nathan Emery, Carla D’Antonio, Christopher Still

**Author notes:** **Corresponding Author**: Nathan Emery.

## Abstract

Across most Mediterranean-type climate regions, seasonal drought desiccates plants, facilitating ignition and the spread of wildfires. Along the California coast, summertime fog has the potential to ameliorate drought conditions and thus reduce plant flammability during a critical time of elevated fire risk. This study investigated the uptake of dry season fog and how it affects live fuel moisture in six dominant shrub species from chaparral and sage scrub plant associations. Fog water uptake was evaluated using stable isotopes of hydrogen and oxygen at several field sites in Santa Barbara County, California. Clear evidence of fog water uptake was identified only in *Baccharis pilularis*, from the sage scrub association. To determine the effects of fog on live fuel moisture, meteorological variables and indices including fog deposition were combined into principal components and the scores regressed against the live fuel moisture loss rate during the summer drought. Fog deposition slowed rates of live fuel moisture loss for all three sage scrub species tested, but it did not affect the chaparral species. Fog is a more regular occurrence in the sage scrub association and thus it is likely that fog ameliorates drought for species that experience consistent fog during the summer months. In coastal California, summer fog can be essential to plant water relations and reduce live fuel moisture loss rates during the summer drought. Understanding these effects is important in the context of changing climate in southern California and Mediterranean-type climate regions around the world.

## Introduction

Fundamental research on the relative importance of different water sources for plants has led to the exploration of the importance of clouds as a direct water source in ecosystems around the world (Weathers 1999). While clouds have indirect effects on plants such as reducing solar insolation and hence evapotranspiration (Burgess and Dawson 2004, Williams et al. 2008, Fischer et al. 2009, Carbone et al. 2013, Baguskas et al. 2014), direct cloud water use in the form of fog uptake can improve water balance (Limm et al. 2009, Baguskas et al. 2016, Emery 2016), enhance carbon assimilation (Berry and Smith 2013, Gotsch et al. 2014) and increase hydraulic function (Laur and Hacke 2014). Fog can also affect community and ecosystem-level processes by changing hydrological cycling through the input of water (Dawson 1998, Matimati 2009, Scholl et al. 2010, Sawaske and Freyburg 2015), thereby altering demography (Baguskas et al. 2016) and affecting range distributions of species (Limm and Dawson 2010). These effects may be pronounced in seasonally dry ecosystems such as the Mediterranean-type climate region of coastal California.

In four of the five Mediterranean regions of the world, including California, wildfire is a natural part of the disturbance regime. While natural fire regimes are influenced by many factors, in southern California extreme fire weather (Davis and Michaelsen 1995, Moritz et al. 2010) and live fuel moisture (Dennison et al. 2008, Dennison and Moritz 2009) are important drivers of wildfire patterns. Live fuel moisture, the ratio of water to dry material in live plants (LFM; Countryman and Dean 1979), is a critical component of plant flammability (Anderson 1970, Martin et al. 1994), indicative of fire intensity (Green 1981) and is incorporated into many fire danger rating systems (Weise et al. 1998, Chuvieco et al. 2004). During the summer drought, LFM generally declines (Miller and Poole 1979) and in southern California this pattern of decline is strongly influenced by spring precipitation (Dennison and Moritz 2009). During the summer drought, fog events occur along the California coast (Leipper 1994, Rastogi et al. 2016), providing shading as well as direct moisture for plants (Williams et al. 2008, Fischer et al. 2009, Baguskas et al. 2014, Emery and Lesage 2015). Fog could potentially alter patterns of LFM decline, yet these effects are unstudied.

While fog could contribute to patterns of LFM, other climatic variables such as rainfall, temperature, relative humidity (which co-varies with fog), wind and solar insolation may also influence the rate of decline in LFM over the summer months. Previous research on the predictors of LFM has investigated seasonal rainfall, drought and fire weather condition indices such as the Keetch-Byram Drought Index (KBDI; Keetch and Byram 1968), Canadian Fire Weather Index (CFWI; Van Wagner 1987) or Fosberg Fire Weather Index (FFWI; Fosberg 1978). The Keetch-Byram Drought Index has been shown to predict LFM of shallow-rooted herbaceous species but not deep-rooted trees in Greece (Dimitrakopoulos and Bemmerzouk 2003). The Drought Code, a component of the CFWI, has been used to predict LFM of shrubs in the Iberian Peninsula (DC; Viegas et al. 2001) and Italy (Pellizzaro et al. 2007). The FFWI was found to predict human-caused ignitions in Switzerland (Reineking et al. 2010) and is a good tool for gauging the influence of weather on wildland fire (Goodrick 2002). It is not unreasonable to expect that the FFWI may predict patterns of LFM as well. While all of these indices incorporate climatic data in various forms, they lack information on fog inundation. During periods of no rain, LFM loss may change depending on cloud shading, fog immersion, and the ability of plants to take up fog water.

For much of coastal California, fog frequency (Leipper 1994, Fischer et al. 2009, Rastogi et al. 2016), vegetation, and fire disturbance patterns (Keeley and Fotheringham 2001) differ by elevation. Chaparral associations tends to grow at higher elevations than the sage scrub association, receive more rainfall, and are dominated by evergreen sclerophyllous shrubs (Hanes 1977) while sage scrub is largely comprised of drought-deciduous shrub species (Westman 1981). While fire frequency for chaparral is highly variable across the state, the natural fire return interval for southern California chaparral is considered to be 30–80 years depending on the location (Franklin et al. 2001, Keeley and Fotheringham 2001, Moritz et al. 2009). At lower and typically drier elevations, sage scrub is the dominant plant association and it is characterized by several drought-deciduous species (Westman 1981) and less frequent fire disturbance (Keeley et al. 2005). While previous studies have demonstrated that several species within sage scrub associations have the potential to take up fog (Cole 2005, Emery 2016), there have been no investigations of how summer fog affects LFM patterns.

This study investigates fog water uptake in coastal Mediterranean California shrub species over two separate summer drought seasons and also how fog deposition and aspects of climate influence LFM in species from two different plant associations. It specifically addresses the following questions:

1. Do coastal California shrub species use fog water during the summer drought, and if so, how does this differ among the species from chaparral and sage scrub associations?
2. How does fog deposition affect LFM trends of these species relative to climatic indices and other meteorological factors during the summer drought period?

Fog frequency along the California coast is inherently variable in space and time (Leipper 1994, Lewis et al. 2003), thus fog water use can be difficult to detect and will be influenced by the quantity of fog deposited as well as plant location within stands. When fog droplets encounter plants, the water adheres to vegetation, drips to the soil surface and wets the upper layers of soil (Fischer and Still 2007, Carbone et al. 2011). We hypothesize that sage shrub species at lower elevations are more likely to encounter low-lying clouds and take up fog water through shallow roots or foliar uptake (Emery 2016). LFM for all species is likely to be influenced by multiple environmental factors; however, lower elevation species are more likely to experience fog events, as fog is typically more common between 200–400m in this region due to the presence of a strong inversion, above which fog formation is unlikely (Fischer et al. 2009).

Thus, we predict that summer fog is more likely to reduce LFM loss in the sage scrub association compared to chaparral and have a stronger influence on plants located adjacent to the coast.

## Methods

### Site selection

This study was conducted at four field sites in Santa Barbara County, CA. The region experiences a Mediterranean-type climate with hot, dry summers and cool, wet winters. The four sites include two along the coast and two in the interior Santa Ynez Valley. The Santa Ynez Valley lies north of the coastal sites and the E-W oriented Santa Ynez Mountains. The field sites were chosen to represent locations of low and high fog frequency, respectively. The sites were selected based on vegetation association, accessibility, and proximity to a meteorological station. The two plant associations are chaparral and sage scrub with one site of each vegetation type in each of the coastal and valley areas.

The coastal field site for sage scrub was at El Capitan State Beach (34°28’N, 120°01’W) in summer of 2011 and nearby Coal Oil Point Reserve for 2012–2013. The shift occurred due to restrictions placed on sampling at El Capitan State Beach. El Capitan State Beach will be referred to as Sage Scrub Coastal One (SSC1, see Figure 1) and is located 16.5 km west of the University of California Santa Barbara. The vegetation there consists of *Baccharis pilularis*, *Artemisia californica* and *Salvia leucophylla.* The soil type is Milpitas-Positas fine sandy loam (El Capitan General Plan, 1979). Coal Oil Point Reserve, (SSC2, Figure 1), is part of the University of California Natural Reserve System and is located two km west of the University of California Santa Barbara campus (34 24’N, 119 52’W). The climate at Coal Oil Point Reserve is very similar to El Capitan State Beach with a mean precipitation of 441±8 mm per year and a monthly mean temperature of 14°C in January and 19°C in August. At Coal Oil Point there are stands of *Artemisia californica* and *Baccharis pilularis*, but not *Salvia leucophylla.* The soil is Concepcion fine sandy loam with intrusions of clay. The chaparral coastal field site (CHC, Figure 1) was located on private property along Painted Cave Road at 780 meters in elevation above the city of Santa Barbara, CA (34 30’N, −119 47’W). According to the nearby San Marcos weather station located 3 km west of CHC at the same elevation, mean January temperature is 12°C, mean August temperature is 22°C and mean annual rainfall is 868±472mm. The chaparral association at this field site is a mesic association with a mix of *Adenostoma fasciculatum*, *Ceanothus megacarpus*, other *Ceanothus* species, and *Arctostaphylos* species. The soil is derived from Coldwater sandstone parent material (Dibblee 1966).

**Figure 1.**
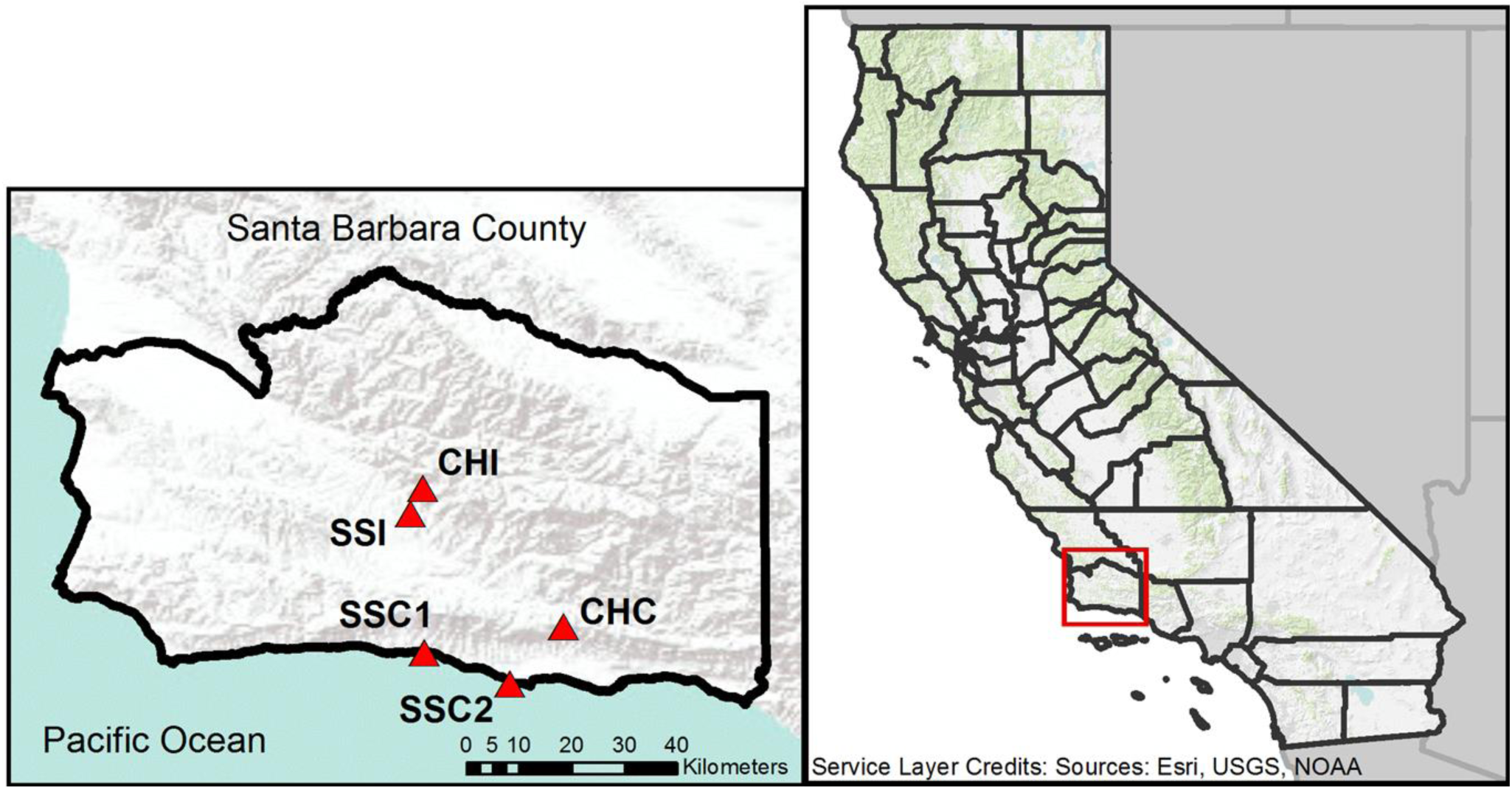
Map of field sites indicated by triangles in Santa Barbara County, California. Sage scrub coastal species were measured at SSC1 in 2011 and SSC2 in 2012–2013.

The sage scrub interior field site (SSI, Figure 1) is located in Sedgwick Reserve, part of the University of California Natural Reserve System, 7.3 km northeast of Los Olivos, CA (34 41’N, 120 2’W) 24 km from the coast. The field site is at 425m elevation with a mean January temperature of 9°C, mean August temperature of 28°C, and mean annual rainfall of 561 mm. SSI is an uplifted Pleistocene alluvial terrace and is mapped as consisting of Ballard Series soils that are generally fine sandy loams or gravelly fine sandy loams (Shipman, 1972). The fog collector and weather station are located adjacent to a mixed stand of *Artemisia californica* and *Salvia leucophylla* (Roberts et al. 2010). Nearby lies a stand of *Baccharis pilularis*, located 0.5 km to the southwest of the weather station and considered part of the SSI field site. The chaparral interior field site (CHI, Figure 1) was located in Sedgwick reserve 4.8 km north of SSI along Figueroa Mountain Road and consisted of mixed *Adenostoma fasciculatum* and *Ceanothus cuneatus.* The soil type is a Toomes-Climara complex. The field site is at 760m elevation and according to the nearby Figueroa Mountain RAWS weather station (991m elevation, 1.9km east of CHI), mean January temperature is 12°C, mean August temperature is 26°C and mean annual rainfall is 544±253mm. For a summary table of field sites, see Table 1.

**Table 1.** Climatic and elevational information for all field sites; chaparral coastal (CHC), sage scrub coastal 1 and 2 (SSC1, SSC2), and sage scrub interior (SSI).

| Table 1 Field Site information |  |  |  |  |
| --- | --- | --- | --- | --- |
| Field Site | Elevation (m) | Mean Annual Precipitation (mm) | Mean January Temperature ($^{\circ}\text{C}$) | Mean August Temperature ($^{\circ}\text{C}$) |
| CHC | 780 | 868 | 12 | 22 |
| CHI | 762 | 544 | 12 | 26 |
| SSC1/SSC2 | 4/10 | 441 | 6 | 24 |
| SSI | 386 | 561 | 10.5 | 19 |

### Species selection

We chose to evaluate dominant species in the sage scrub association that differ in phenology and rooting depths. *Artemisia californica* is a drought-deciduous shrub considered to have a shallow root system (Kirkpatrick and Hutchinson 1977) and may use fog water in the late summer (Emery and Lesage 2015). *Salvia leucophylla* is also a shallow-rooted drought-deciduous species with broader leaves than *A. californica*, and a patchier distribution (Kirkpatrick and Hutchinson 1977). By contrast, *Baccharis pilularis* is a deep-rooted evergreen shrub found throughout California (Wright 1928, Ackerly et al. 2002) that can be abundant in the coastal zone. *Artemisia californica* and *B. pilularis* were measured from 2011–2013 while *S. leucophylla* was measured only for the 2011 field season, as SSC2 lacked *S. leucophylla.*

Chaparral is typified by evergreen sclerophyllous-leaved shrub species (Hanes 1977). *Adenostoma fasciculatum* is a widespread evergreen shrub and has small needle-like leaves (Kummerow et al. 1977). This species can have variable rooting depth (Kummerow et al. 1977) and is the main species measured by the US Forest Service to monitor live fuel moisture as an index of fire danger in chaparral (Burgan 1979). This study also included two species of the genus *Ceanothus*, *C. megacarpus* for the coastal chaparral site and *C. cuneatus* for the interior chaparral site. Both species are in the subgenus *Cerastes*, a more drought-tolerant subgenus of *Ceanthous* (Davis et al. 1999) with species known to have shallow roots (Kummerow et al. 1977). Of all *Ceanothus* species, *C. megacarpus* and *C. cuneatus* tend to occupy areas of high drought severity (Meentemeyer and Moody 2002) and are resistant to water stress-induced cavitation (Davis et al. 1999). *Ceanothus megacarpus*, in particular, is a highly drought tolerant evergreen shrub species (Gill and Mahall 1986, Jacobsen et al. 2007).

### Water isotopes and tracking of fog uptake

This study used stable isotopes of hydrogen and oxygen to identify fog water uptake into stem tissue (Dawson et al. 2002). Isotopic analyses were conducted at each field site except for CHI as no fog deposition was detected there for the duration of the study. At each of the other field sites (SSC1, SSC2, SSI and CHC), groundwater, rainwater and fog water samples were collected for three years (2011–2013) and analyzed for their stable isotopic ratios. Groundwater was collected from drilled wells within one kilometer of each field site. Rainwater for the SSC1 and SSC2 field sites was collected at SSC2 (Coal Oil Point Reserve). Rainwater was also collected at CHC and SSI and brought to the lab after every rain event. A Nalgene container with several centimeters of mineral oil was used to trap rainwater and prevent evaporative fractionation for subsequent isotopic analysis. For all field sites, fog deposition was measured using a fog collector modified from Fischer et al. (2007) containing a Nalgene with several centimeters of mineral oil. The container was swapped out approximately every two weeks in the field and brought to the lab to measure volume, extract the water and filter using a 0.45 μm cellulose acetate filter. Fog collectors were not permitted at SSC1, so fog water data from SSC2 was used for SSC1. Fog, rain and groundwater samples were analyzed for stable isotopes of hydrogen and oxygen using Isotope Ratio Infrared Spectroscopy at the Stable Isotope Biogeochemistry Lab at UC Berkeley. Fog and rainwater collection occurred at SSC2, CHC and SSI for all three years of the study. Fog collection occurred during the summer months, from May to the first rain event of the fall. Rain water isotopic values were used to develop a local meteoric water line for the Santa Barbara region.

In semi-arid environments, water isotopes in the upper soil profiles experience evaporative fractionation. As a result, the isotopic ratio of water taken up by roots is generally more enriched in the heavier isotope than the source water, thus making it difficult to attribute proportional water source use in plants. Using the method outlined by Corbin et al. (2005), we developed evaporative correction regressions for sage scrub and chaparral plant associations. This consisted of taking soil water isotope samples from the soil at 0–2 cm and 2–5 cm depth at one-hour intervals on a sunny day from sunrise to midday. Samples were placed in 20ml scintillation vials, which were then sealed with parafilm and frozen for future extraction. Sampling across this time interval should capture evaporative fractionation of surface soil water isotopes and captures how fog or rainwater may fractionate at the soil surface. Four soil sampling events at SSC2 (4/26/2013, 7/10/2013, 8/26/2013, 4/5/2014) and two at CHC (6/7/2013, 8/12/2013) were taken over the summers of 2013 and 2014 to develop regression lines for the two plant associations. The slopes of both depths were similar within sites so the data were aggregated by site to develop evaporative corrections. These association-specific regression lines were used to correct plant water samples for all field sites to the local meteoric water line (see supplementary material, Appendix I). The SSC2 correction was used for sage scrub sites SSC1, SSC2 and SSI as these sites consisted of similar fine sandy loam soils. The CHC correction was used for the chaparral site CHC. In order to propagate error associated with the evaporative corrections, we included the 95% confidence interval of the slope for each regression along with the mean slope when running a mixing model for plant water samples.

To develop a reference soil oxygen isotope profile, soil water samples were collected at two locations within the CHC and SSC2 field sites on 7/5/2012 and 9/12/2012. At midday, five soil samples were collected at 0–5 cm, 5–10 cm, 10–20 cm and 30–50 cm in depth. The soil was placed in a 20ml scintillation vial, the lid sealed with parafilm and immediately frozen for future extraction (see supplementary material, Appendix I). Soil water isotopes can vary widely across sampling locations and time of year (Gazis and Feng 2004). The profile developed for this study is not meant to be an extensive record nor is it corrected for evaporative fractionation, but act as a reference for interpretation of isotope ratios at these field sites.

Plant stem water samples were collected monthly in 2011 and twice monthly in 2012 and 2013 in order to ensure capture of samples from before and after measurable fog events. Sampling consisted of clipping a 10cm section of suberized stem from an individual shrub, quickly removing the bark, placing in a 20ml scintillation vial, placing parafilm over the lid and immediately freezing for future extraction. Stem samples were collected from the same three individuals per species per site on each date of LFM collection for three years.

### Fog detection in stem water

To evaluate whether species and individuals could use fog, the authors reviewed the fog deposition record at a given field site and selected stem samples that had been taken prior to and after a significant period of fog deposition. To contain costs, only one date following the largest deposition of fog was selected for isotopic analysis for each field site. As no measurable fog was collected at CHI, no stem samples were analyzed for this site. Preference was given to late summer fog events when spring rainfall was less likely to influence the water budget of a given shrub individual (see Table 2). All stem and soil samples were extracted using cryogenic vacuum extraction (Ehleringer et al. 2000) in a laboratory at UC Santa Barbara and the Stable Isotope Biogeochemistry Lab at UC Berkeley. The stem and soil water samples were then analyzed for hydrogen and oxygen isotopes on an Isotope Ratio Mass Spectrometer (model Delta plus XL; Finnigan MAT, Breman, Germany) at the Stable Isotope Biogeochemistry Lab at UC Berkeley.

**Table 2.** Dates selected for stem water isotopic analysis. Plant samples were selected prior to and after a significant fog event at each field site (except for CHI).

| Table 2 Stem isotope sampling dates |  |  |  |  |  |
| --- | --- | --- | --- | --- | --- |
| Field Site | Species | Date of sampling |  | Fog collection (ml) | Date of last rainfall (>6mm) |
|  |  | Pre-fog | Post-fog |  |  |
| CHC | *A. fasciculatum*,<br>*C. megacarpus* | 5/3/2013 | 5/17/2013 | 245.7 | 5/6/2013 |
| SSC1 | *A. californica*,<br>*B. pilularis*,<br>*S. leucophylla* | 8/30/2011 | 9/29/2011 | 217.2 | 6/6/2011 |
| SSC2 | *A. californica*,<br>*B. pilularis* | 8/12/2013 | 9/3/2013 | 94.5 | 4/1/2013 |
| SSI | *A. californica*,<br>*B. pilularis*,<br>*S. leucophylla* | 8/1/2011 | 8/30/2011 | 193.97 | 6/6/2011 |

We used the MixSIAR package v3.1 in R (Stock and Semmens 2013) to determine proportional fog water uptake in five shrub species (No fog was detected at the field site with *C. cuneatus*). The hierarchical Bayesian mixing model incorporates uncertainty associated with the isotopic ratios of water sources and estimates the proportional use of a given water source. Stem water isotopes were generally more enriched than source isotopes (Figures 2a-c) and needed to be corrected for evaporative fractionation (Corbin et al. 2005). When entering an individual plant’s stem water isotopic ratio into the model, we included the 95% confidence interval of the slope associated with the corrected stem water samples. Each individual plant sample was represented by 3 isotopic ratios, the mean corrected value and the two 95% confidence interval bounds (See Figure 2D). This conservative approach to determining the proportion of fog water in stem tissue propagates error and uncertainty associated with correcting for evaporative fractionation in arid soils. Following the MixSIAR manual (Stock and Semmens 2013), we ran the hierarchical Bayesian mixing model for each field site, as they had different water sources. Discrimination values for the model were set to zero as there is no isotope fractionation during plant water uptake (Dawson and Ehleringer, 1991; Ehleringer and Dawson, 1992). The Monte Carlo Markov Chain parameter was set to “normal” with a chain length of 100,000 and a burn-in of 50,000. Additionally, we used the “Process only” error structure. All statistical analyses were accomplished using R version 3.3.0 (R Core Development Team, 2016).

**Figure 2(A-D).**
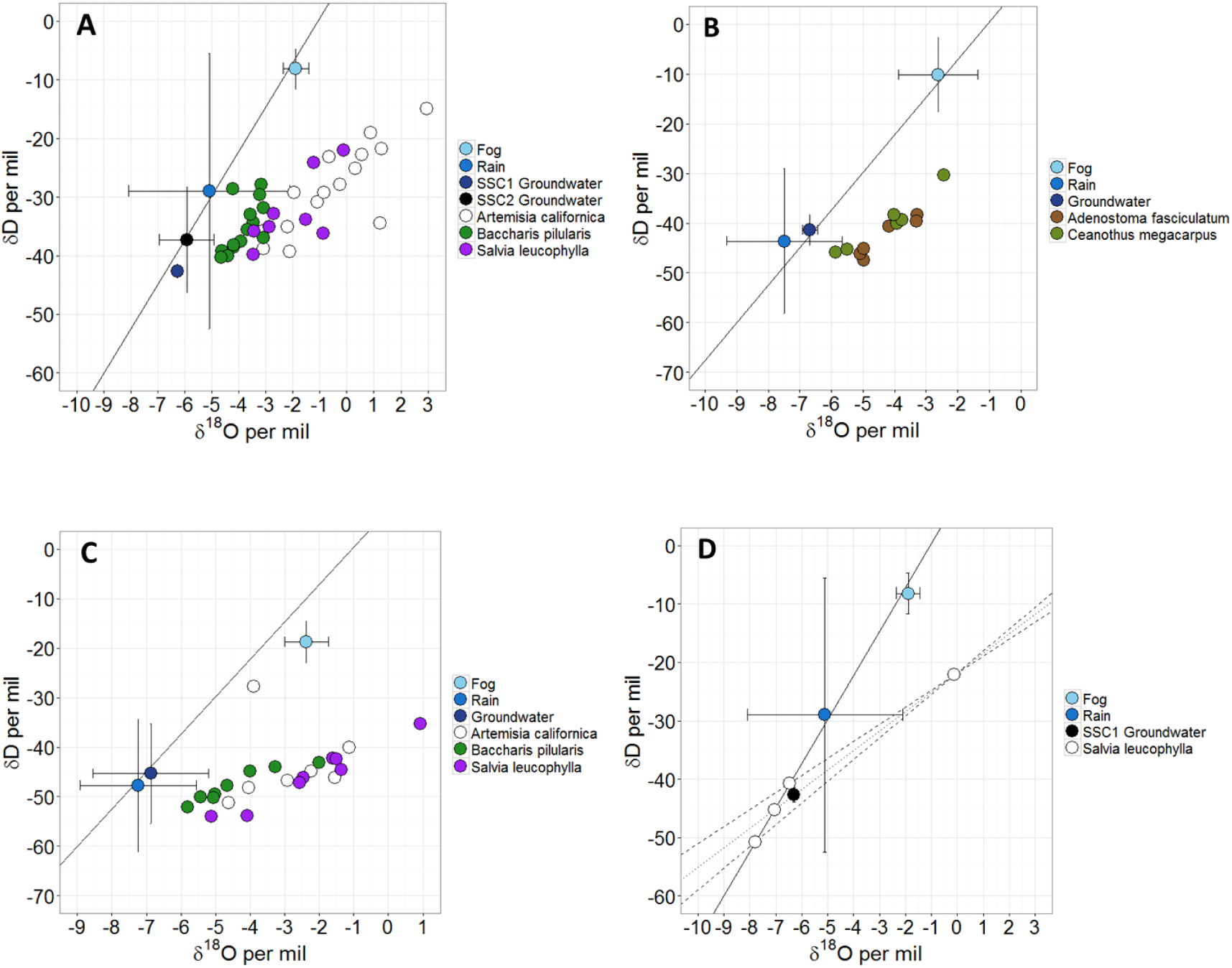
Isotopic ratios of water sources and plant water samples for SSC1/SSC2 (A), CHC (B), and SSI (C). Each water source represents the volume-weighted average value from 2011–2013 (with standard error bars). The line represents the local meteoric water line (LMWL, slope of 7.57). Plant water samples fall to the right of the LMWL. An example of correcting for soil water evaporation at SSC1 and the 95% confidence intervals of the slope (D).

### Live fuel moisture sampling

The standard US Forest Service procedure for sampling LFM involves collecting plant material at 2:00 PM on dry days, with a max stem diameter of 1/8^th^ inch and removing dead material and reproductive tissue. It also includes the separation of plant material into new and old growth (Countryman and Dean 1979). For certain species in this study, particularly *B. pilularis*, it was impractical to remove all reproductive tissue and for others it was difficult to consistently differentiate between old and new plant material. In order to compare LFM across all six species in a repeatable way, a method was developed for LFM sampling that integrated new and old plant material present at any given time point. In the field, two south-facing (when possible) live branches were randomly selected and 15 cm of terminal stem were removed and placed in a water-tight Nalgene container. No more than 30% of the sample was reproductive tissue and all material along the 15 cm was included except for visibly dead material, which was removed. Due to travel limitations among field sites, sampling was not simultaneous and occurred between 1:00 PM and 5:00 PM on fog-free days. Once collected, samples were brought back to the lab at the University of California Santa Barbara. They were wet weighed, oven dried at 80°C for at least 48 hours and reweighed. LFM was calculated as:

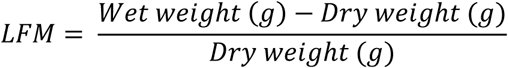

LFM measurements began after what was anticipated to be the last spring rainfall and ended with the first (>2mm) rain event of the fall season. For the 2011 field season, LFM was measured once a month for 12 individual plants per species per site. Measurements began in April and continued until November 2011. For 2012 and 2013, LFM was measured twice a month for 10 individuals per species per site. Several of these individuals at each field site were the same individuals sampled for fog water uptake analysis. Measurements for 2012 began in July and went until September, while in 2013, sampling started in May and went until September.

Because this study used a different LFM sampling method than that of the US Forest Service, the authors compared the two methods for *A. fasciculatum*. This species is the most commonly measured species in Californian USFS inventories. In June, July and August of 2014 we measured ten individual shrubs at the CHC field site using our method, as well as the USFS method of separating new LFM and old LFM (Countryman and Dean 1979). The LFM values obtained by our method are very similar to the average of old and new LFM (see supplementary material, Appendix II). A linear regression between the average of new and old LFM (USFS) and the method used in this study yields a linear relationship with a slope of 0.91 and an R^2^ of 0.92 (intercept = −2.97). This suggests that our method accurately reflects the LFM of *A. fasciculatum* across the summer season. However, other species in this study vary widely in physiology and reproductive phenology. For several species, it was difficult to consistently remove reproductive material and differentiate between old and new material, as prescribed by the USFS. The authors decided to use a standard method across species to enable comparison of LFM values. While this method is different from the USFS method, it is more general in that it quantifies the relative water content of live vegetation for multiple dominant shrub species in California coastal plant associations.

### Predicting LFM using meteorological factors and indices

This study used meteorological data from nearby weather stations to characterize climatic conditions and calculate several drought and fire behavior indices for analysis of LFM data. Meteorological information was collected from weather stations at SSC2 (used for SSC1 and SSC2), CHC, and SSI (Roberts et al. 2010). From 2011 until June of 2012, meteorological data for CHC came from the San Marcos Pass weather station (ID: KCASANTA45), 2.5km west of CHC at 600m in elevation. In June of 2012, researchers from UC Santa Barbara installed a meteorological weather station at CHC, and data was collected at the CHC site from this date on (values from nearby RAWS station KCASANTA45 and this new station follow a 1:1 relationships). Environmental conditions at CHI were measured at the nearby Figueroa Mountain RAWS weather station (991m elevation, 1.9km east of CHI). Meteorological data used in the analysis of all field sites included temperature, relative humidity, wind, rainfall, vapor pressure deficit and dew point depression (Summarized information is available in supplementary materials Appendix III). In addition to meteorological data, this study also quantified fog deposition using fog collectors modified from Fischer et al. (2007) with Nalgene containers to collect fog water. Fog water was collected during the summer months approximately every two weeks. It was difficult to determine when fog deposition occurred within a sampling period so the volume of fog water collected was divided by the number of days elapsed between two sampling dates. This metric, fog per day, represents the average daily fog deposition between any two sampling dates.

Several indices were calculated using meteorological data to model drought, soil moisture and fire weather. The Keetch-Byram Drought Index, Fosberg Fire Weather Index, Drought Code, and Duff Moisture Code (Appendix III) were derived using the open-access Fire Danger Index Functions (http://www.atriplex.info/index.php/Fire_Danger_Index_Functions_in_R). All environmental factors and indices were averaged over two-week intervals prior to a given LFM sampling date, except for rainfall which was summed for the two-week period. Rain was treated differently than fog deposition as rain likely accumulates in the soil profile over a given time period. Two weeks was the minimal LFM sampling interval and as daily weather conditions are unlikely to affect LFM, it was important to aggregate over a period of time preceding a LFM measurement. (see supplementary material, Appendix III).

We used R version 3.3.0 for all analyses (R Development Core Team 2014). Analysis of environmental variables was conducted for the CHI field site separately because no fog was measured at this field site. Because many of the aggregated indices and meteorological factors were correlated, we used the corrplot package to identify and remove redundant variables and the prcomp() function from the stats package in R to perform a principal components analysis (PCA) of the variables. Principal Component Analysis is a standard statistical method used to reduce intercorrelated variables (in this case meteorological ones) into a few uncorrelated components that explain much of the variance (Rencher 1998). For sites SSC1, SSC2, CHC, and SSI, variables selected for the PCA included the Keetch-Byram Drought Index (KBDI), Duff Moisture Code (DMC), Drought Code (DC), Fosberg Fire Weather Index (FFWI), afternoon vapor pressure deficit (VPD), nighttime Dewpoint Depression (DptDep), Fog per day, Temperature, Wind, and Rain. For the CHI field site, variables selected for the PCA included the KBDI, DMC, DC, FFWI, afternoon VPD, nighttime DptDep, and Rain. For both analyses, all variables were log-transformed and centered to control for different scales prior to undergoing PCA. These principal components were then used to predict changes in LFM loss during the summer drought.

### Live fuel moisture analysis

This study explored environmental factors that affect the change in LFM over a time interval because a single LFM measurement does not reflect the direction of moisture loss or gain. An individual shrub may have decreasing or increasing LFM when a measurement is taken. Whether the change in LFM is positive or negative may be directly related to antecedent climatic conditions. In order to understand the change in LFM from one date to the next, we modeled the proportional rate of change in LFM from time point 1 to time point 2. The resulting LFM rate was the following:

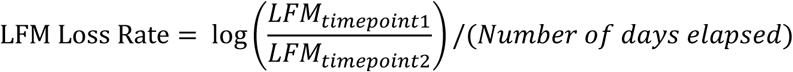

Environmental Influences on LFM: Linear Mixed Effects Model

We chose to use a linear mixed effects model to determine the relative influence of environmental factors, indices, and fog deposition on the LFM loss rate for each shrub species. LFM loss rate was analyzed by species, including all field sites where those species were found. This means, for example, that *C. cuneatus* was only evaluated at CHI, while *B. pilularis* was evaluated across SSC1, SSC2 and SSI. All years of LFM data were used in the analysis. For each individual plant, the initial LFM at the start of each summer was designated a fixed effect as the LFM loss during the summer is affected by how well hydrated the plants are at the beginning of the summer. Additionally, “shrub individual” was designated as a random effect as individual shrubs at the same field site tended to follow a similar pattern of LFM change across each summer season. For species that were measured at multiple sites, field site was also set as a random effect in the model.

For *A. californica*, *A. fasciculatum*, *B. pilularis*, *C. megacarpus*, and *S. leucophylla*, four principal components (from the general PCA) were used in the linear mixed effects model to predict LFM loss rate using the maximum likelihood method from the nlme package in R. For *C. cuneatus*, three principal components from the CHI-specific PCA were used. Using Akaike’s Information Criterion (AIC), models of LFM loss rate were ranked. When the model for PC3 (representing fog and rain) had the lowest AIC, except for *C. cuneatus*, the significance of fog and rain was evaluated separately using the linear mixed effects and the restricted maximum likelihood approach to determine significance of fog or rain on the LFM loss rate for a particular species.

## Results

### Fog Collection

Fog deposition across years was highly variable and differed among field sites (Figure 4). In 2011 there was fog at both CHC and SSC2 in the early summer (May/June) as well as in the late summer at SSC2 (August/September). For all three years at the interior field site, SSI, fog deposition was low throughout the summer months and the water collected likely included dew. No fog water was collected at CHI for all three summers of the study.

### Water sources and their isotopic signatures

As expected, fog water was more enriched in the heavier isotopes of hydrogen and oxygen compared to groundwater and rain for the three sites that experienced fog deposition, sage scrub coastal, chaparral coastal, and sage scrub interior (SSC1/SSC2, CHC, SSI; Figures 2A-C). This is consistent with previous research on isotopic ratio differences in source water (Scholl et al. 2010). The isotopic ratio of fog water was similar between the two coastal sites (CHC, SSC2) and slightly depleted in the hydrogen isotope for the SSI site. At the sage scrub coastal sites, rain was variable and slightly enriched compared to groundwater for both sites (SSC1 and SSC2, Figure 2A). At the chaparral coastal site (CHC, Figure 2B) and sage scrub interior site (SSI, Figure 2C), rain was slightly more depleted than groundwater. The local meteoric water line (LMWL), y = 7.5684x + 8.0852, was developed from three years of rainwater isotopes from all field sites. Plant water samples for all species were more enriched in the hydrogen and oxygen isotopes than the water sources and fell to the right of the LMWL (Figures 2A-C). These patterns reflect findings from nearby Santa Cruz Island (Fischer et al. 2016).

For both CHC and SSC2, the oxygen isotopic ratio was depleted for the top layer of soil (see supplementary material, Appendix I). Subsequent depths were initially more enriched near the top layer and became more depleted with depth. These patterns are typical of arid, low rainfall climates (Hsieh et al. 1998). For all soil depths, the oxygen isotopic ratio was more enriched in the heavier isotope for SSC2 than for CHC.

The linear regression used to correct for soil water evaporative fractionation differed by field site (see supplementary material, Appendix I). Consistent with the soil oxygen isotopic profile, soil water was generally more enriched in both isotopes at SSC2 compared to CHC. The slope of the regression at CHC was 2.76 and 3.33 at SSC2. An example of correcting for soil water evaporative fractionation is shown in Figure 2D.

### Fog water uptake

The MixSIAR model reveals that most individuals of all five species of shrubs did not use fog water in the interval over which their tissues were collected (Figure 5A-D). The only species that had noteworthy proportions of fog, *B. pilularis*, differed in fog proportion by shrub individual (identified by shape in Figure 5B, C). At SSI in August of 2011, several *B. pilularis* individuals consistently had ~10% fog water in their stem tissue (Figure 5B). At SSC1 in September of 2011, one individual *B. pilularis* shrub went from 0–5% fog to ~30% fog over the course of one month, while another increased from 0–5% to ~15% fog (Figure 5C).

### Live fuel moisture analysis

The patterns of LFM loss during the summer drought period followed an exponential decay function (Figure 3). Most study species lost water rapidly in the early summer and approached an asymptote by mid-summer. Unlike most of the species measured, *B. pilularis* maintained relatively high LFM throughout the late summer months.

**Figure 3.**
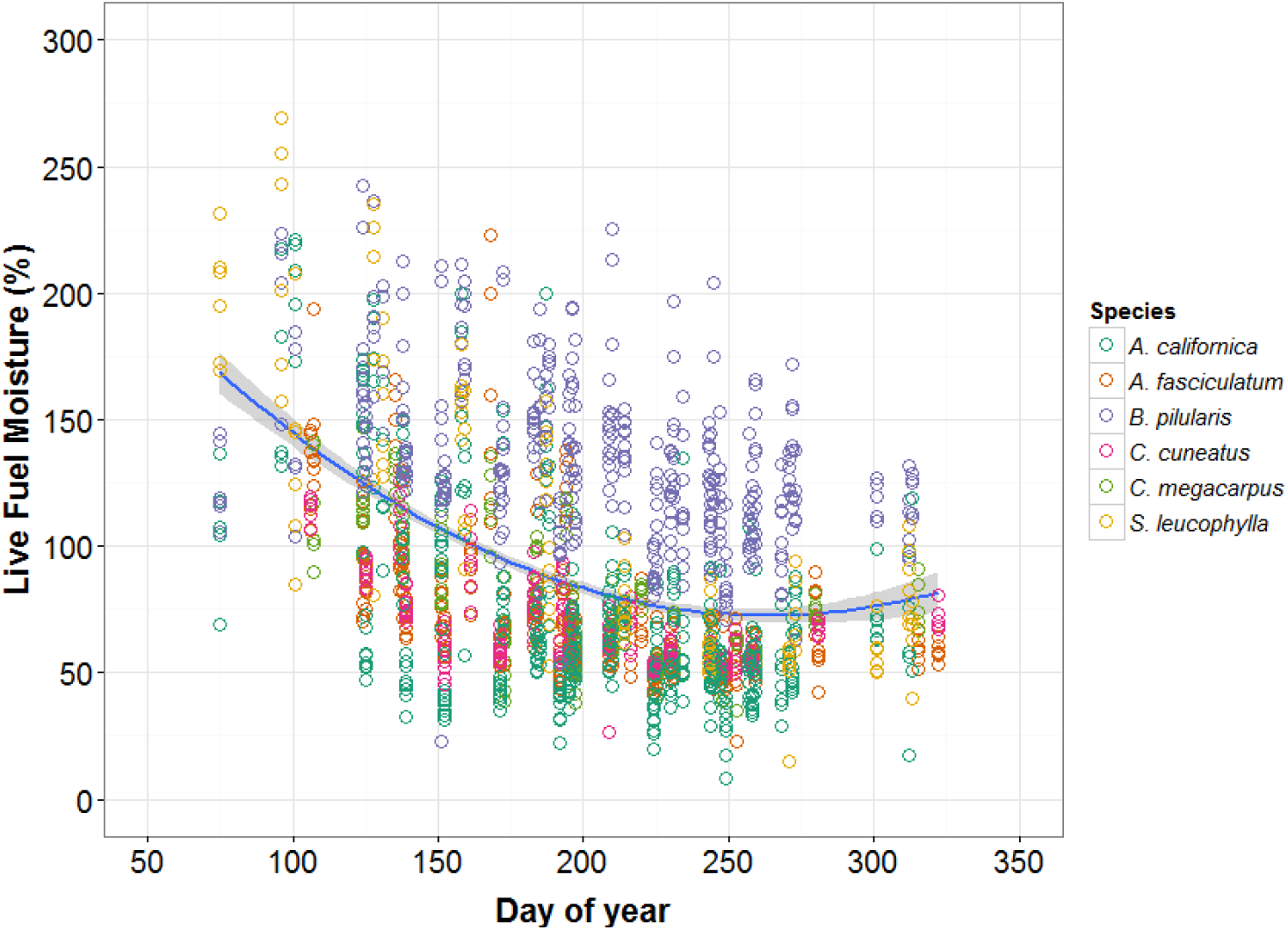
Live fuel moisture of all species over three summers (2011–2013) fitted with an exponential decay regression line and 95% confidence intervals. Each color represents a different species.

The principal component analyses for SSC1, SSC2, CHC, SSI yielded four principal components that explained 87.7% of the variance. The explanations of principal components were based on factors with eigenvalues greater than 0.4 or less than −0.4. Principal Component I (PC I) represents shallow soil moisture deficit and was primarily composed of the Keetch-Byram Drought Index, Duff Moisture Code and temperature (see loadings in Appendix IV). PC II represents evaporative strength of the atmosphere and was composed mainly by the Fosberg Fire Weather Index and afternoon VPD. PC III is moisture availability and was primarily fog and rain while PC IV was deeper soil moisture availability and composed of the Drought Code and rain. The CHI-specific analysis yielded three principal components that explained 91.3% of the variance. PC I is a combination of factors including rain, afternoon VPD, KBDI, and nighttime DeptDep (see loadings in Appendix V). PC II represents fire weather conditions and soil moisture and is composed of FFWI and DC. PC III represents shallow soil moisture and is mainly DMC and nighttime DeptDep. The principal components for the general analysis and CHI-specific analysis were used in the subsequent linear mixed effect models.

The results of the linear mixed-effects model selection differed primarily by plant association (Table 3). In chaparral, the best model for predicting the LFM loss rate in *A. fasciculatum* and *C. megacarpus* was PC IV representing deep soil moisture and rainfall. At the CHI field site, the LFM loss rate of *C. cuneatus* was best predicted by PC II representing fire weather conditions and deep soil moisture.

**Table 3.** Results of linear mixed effects model. Model selection based on AICc of principal components.

| Species | Number of observations | Field sites | Model AIC |  |  |  | Model log-likelihood |  |  |
| --- | --- | --- | --- | --- | --- | --- | --- | --- | --- |
|  |  |  | PC1 | PC2 | PC3 | PC4 | PC1 | PC2 | PC3 |
| *A. californica* | 324 | SSC1, SSC2, SSI | -1733.15 | -1732.88 | -1749.92* | -1733.15 | 872.57 | 872.44 | 880.96 |
| *A. fasciculatum* | 132 | CHC | -813.60 | -816.46 | -822.28 | -835.19* | 411.80 | 413.23 | 416.14 |
| *C. megacarpus* | 133 | CHC | -885.61 | -888.90 | -881.46 | -891.77* | 447.81 | 449.45 | 445.73 |
| *B. pilularis* | 329 | SSC1, SSC2, SSI | -1958.71 | -1956.34 | -1962.43* | -1955.98 | 985.36 | 984.17 | 987.21 |
| *S. leucophylla* | 72 | SSC1, SSI | -436.64 | -436.38 | -441.14* | -437.76 | 224.32 | 224.19 | 226.57 |
| *C. cuneatus* % | 133 | CHI | -885.61 | -888.90* | -881.46 | n/a | 447.81 | 449.45 | 445.73 |

\* indicates best model
% model does not include fog deposition

In the sage scrub association, the LFM loss rate of all three species, even the deep-rooted *B. pilularis*, was best predicted by PC III which represents fog deposition and rainfall. When analyzed separately, rainfall and fog differed in significance. Fog significantly affected LFM loss rate in *A. californica* (p=0.001), *S. leucophylla* (p<0.001), and *B. pilularis* (p=0.013). However, rainfall effects on the LFM loss rate were only significant for *A. californica* (p=0.012).

## Discussion

While fog deposition varied across sites and years, late summer fog deposition tended to occur at lower elevations. *Baccharis pilularis* was the only species that showed isotopic evidence of fog water uptake, demonstrated at two separate field sites (Figure 5A-D). Despite lack of fog uptake across species, summer fog reduces the rate of LFM loss for the three sage scrub species (Table 3). LFM decline in chaparral species, however, was not significantly influenced by fog but was influenced by soil moisture and fire weather conditions. The effects of direct fog water uptake are likely ephemeral, but long-term exposure to summer fog can significantly influence LFM for species in lower elevation plant communities.

### Fog patterns and fog water uptake

Fog deposition was highly variable among field sites and years (Figure 4). Fog tends to occur along the coast in the early summer at both low (sage scrub coastal) and high elevations (chaparral coastal). However, late summer fog primarily occurred at lower elevations as the inversion strengthened and lowered. Fog deposition at the nearby Channel Islands reflects a similar pattern with summer clouds remaining below 400m in elevation (Fischer et al. 2009). This suggests that chaparral species at higher elevations do not experience summer fog as frequently and are thus less likely to be influenced by this water source during the late part of the summer drought. These the patterns of cloud-base height along the California coast may change in the future as cloud levels rise from urbanization (Williams et al. 2015) or changing climatic patterns. Urban development at lower elevations may reduce fog inundation for the sage scrub association if nighttime warming occurs from the urban heat island effect (Oke 1982) and reduces the likelihood of fog formation. In the Santa Ynez Valley, at the SSI site, fog deposition was low but consistent throughout the summer (Figure 4). This could reflect early morning dew that registers as fog, although early morning fog is known to occur in the Santa Ynez river basin (Upson and Thomasson 1951). Fog deposition patterns, while variable, may change in the future particularly if upwelling patterns change as increased upwelling increases the marine cloud layer along the California coast (Bakun 1990, Snyder et al. 2003). These changes could influence plant water relations and subsequently affect species composition of lower elevation sage scrub communities.

**Figure 4.**
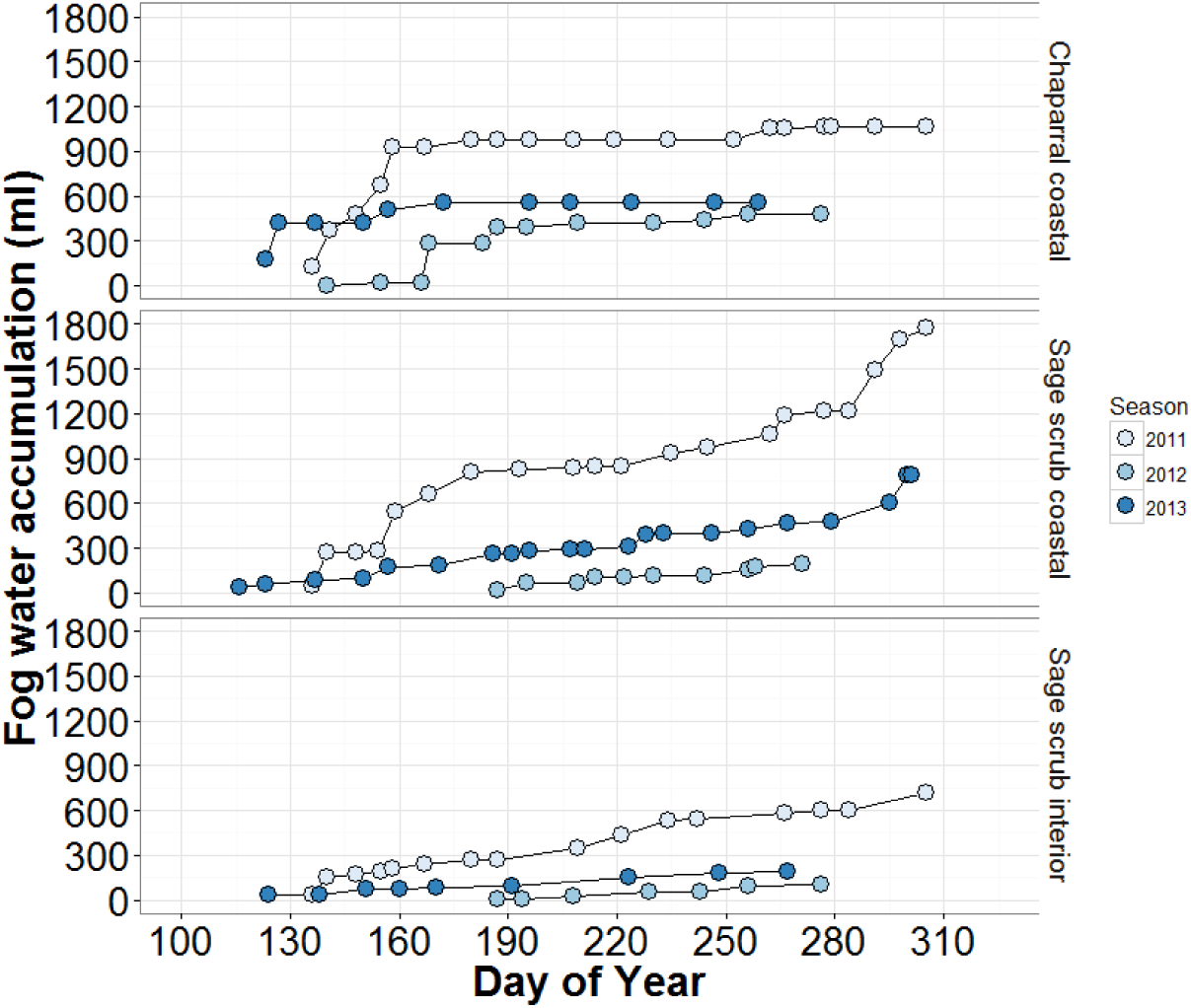
Fog deposition at CHC, SSC2 and SSI for the summers of 2011–2013. These data are the fog water accumulation over the summer months for a given year.

The only species that demonstrated fog water uptake was *Baccharis pilularis*, a deep-rooted evergreen shrub found at lower elevations (Figure B-C). Although fog water uptake was not detected in four of the shrub species sampled, this does not mean conclusively that these species cannot take up fog. *Adenostoma fasciculatum*, *Artemisia californica*, and *Salvia leucophylla* can take up fog water through their leaves in controlled settings (Emery 2016). The time elapsed between sampling dates (two weeks to one month), may have prevented detection of fog in the stem tissue of some species. Fog uptake in shrubs is possible, but the residence time of fog water may be short. The additional water in plant tissue during the summer drought may quickly transpire and leave no isotopic evidence. Lastly, this study constructed the local meteoric water line from rain water isotopes assuming that soil water is primarily derived from rain. Fog water is more enriched in heavier isotopes and falls off the line (Figures 2A, 2C). It is possible that this study underestimated fog uptake as the evaporative fractionation correction does not account for the enriched source water. However, if this was the case, the underestimation would have had little to no effect on the isotope analysis.

Variable fog frequency suggests that any fog water uptake would be an opportunistic event for many plants and is not frequent enough or in high enough quantity to influence all shrub individuals for a given plant association. Fog water uptake can be highly variable among species as observed in northern Californian plant associations (Dawson 1998, Corbin et al. 2005). In chaparral, *A. fasciculatum* and *C. megacarpus* did not use fog water in the early summer months (Figure 5A). Prior to sampling, there was a rain event which may have provided rain water to deeper roots (Table 2). Fog drip usually wets only the litter layer and top few centimeters of soil (Ingraham and Matthews 1995, Dawson 1998, Corbin et al. 2005, Fischer and Still 2007, Carbone et al. 2013, Fischer et al. 2016). If chaparral shrubs have roots primarily distributed at greater depths, they are unlikely to access fog drip. *Adenostoma fasciculatum* is reported to have a variable rooting system while *Ceanothus* species tend to have relatively shallow roots (Hellmers et al. 1955, Kummerow et al. 1977, Kummerow et al. 1978). Most shrub canopies in the chaparral study area are very tall (>2 meter), and cover the entire soil with two meters of stems and leaves. We observed that fog events during the study rarely penetrated the canopy to reach the ground beneath these dense canopies.

**Figure 5.**
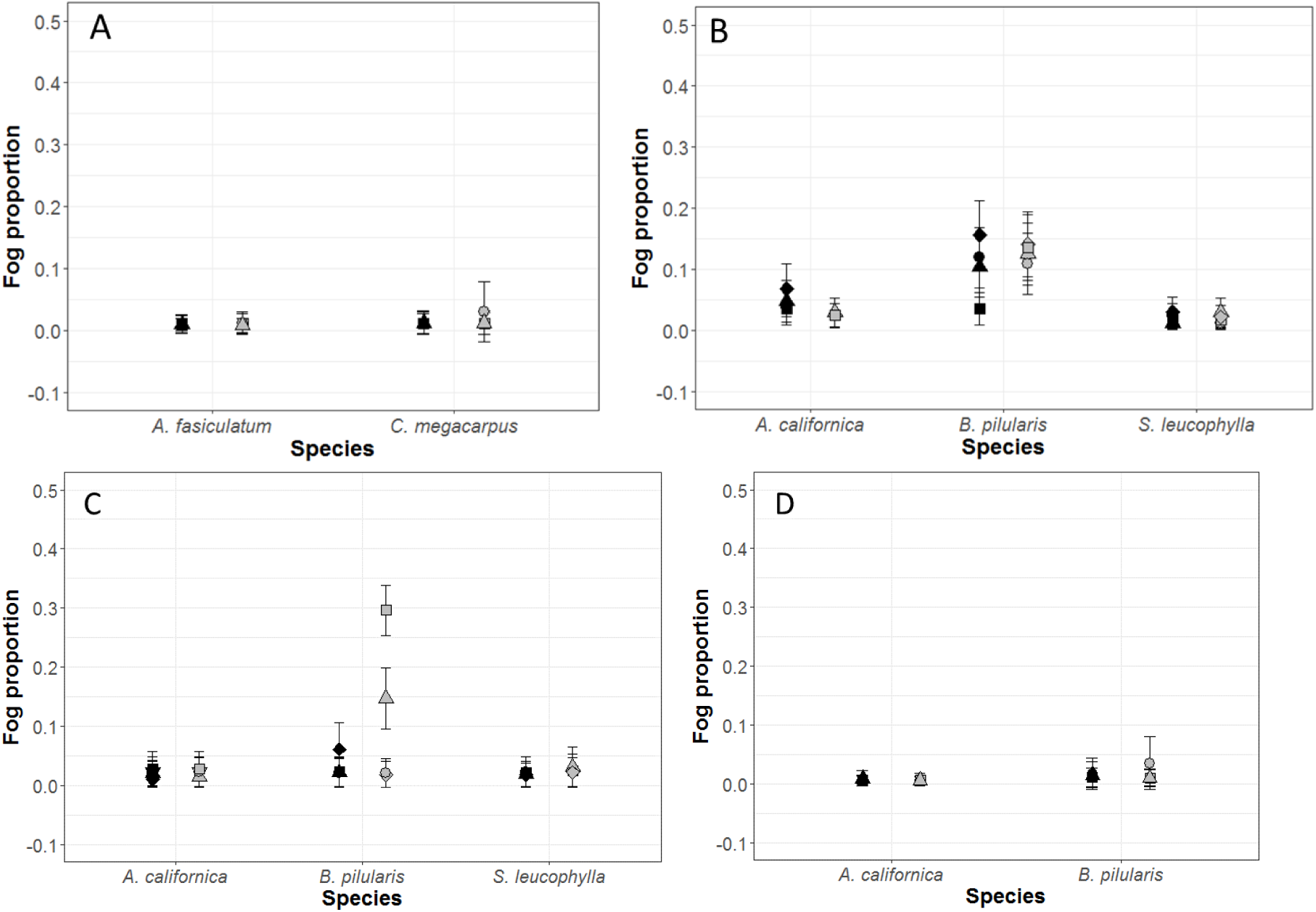
Proportional fog water uptake for all field sites. Pre-fog event samples are in black, post-fog in gray. Each shape represents the identification of a single shrub individual. The points and lines represent the mean and standard deviation output from the MixSIAR mixing model. Note the adjusted y-axis for CHC (A), SSI (B), SSC1 (C), and SSC2 (D).

In the sage scrub association, although all three study species can undergo foliar uptake of fog (Emery 2016), *B. pilularis* was the only species that showed fog water uptake during the late summer (Figure 5B-C). The two drought-deciduous species, *A. californica* and *S. leucophylla*, have shallow roots (Kirkpatrick and Hutchinson 1977) and are known to use small rain events (Gray 1982). However, these species drop many of their leaves by the late summer and may be physiologically dormant. Direct fog uptake may occur, but the effects are likely short-lived. Despite the lack of isotopic evidence of fog water in plant tissue, new leaf growth has been observed in *A. californica* following late summer fog events and prior to the first fall rain events (Emery, *personal observations*). Although *B. pilularis* can take up fog water through leaves (Emery 2016), root uptake is also possible if fog deposition is heavy. At two of the field sites, *B. piluaris* was using non-trivial proportions of fog water (Figure 5B-C). Even though *B. piluaris* is thought to have deep roots that tap into groundwater (Wright 1928, Ackerly et al. 2002), established stands can still have roots present in the top 15cm of soil (Zavaleta and Kettley 2006) and potentially access fog drip in the summer months. Fog water may be important for young *B. pilularis* to establish (Kidder 2015) and act as a supplementary water source as the plants’ rooting distribution becomes deeper with age (Zavaleta and Kettley 2006).

### Fog and live fuel moisture

Chaparral is a widespread fire-prone plant association in California (Epling and Lewis 1942, Hanes 1971). To understand fire risk and how fire spreads through ecosystems, it is important to consider plant flammability (Anderson 1970, Gill and Zylstra 2005) and LFM is a critical component of plant flammability (Anderson 1970, Martin et al. 1994). Although the chaparral shrub species in this study took up fog early in the summer drought, fog did not affect the rate of LFM loss. Instead, results from our analyses suggest that soil moisture (DC) and rainfall are important contributors to LFM decline in *A. fasciculatum*, arguably the dominant species across much of coastal California chaparral (Table 3). Previous research has shown that March-May rainfall is critical to the summer LFM decline in *A. fasciculatum* (Dennison and Moritz 2009). These rain events are typically the last precipitation inputs before the summer drought and influence soil moisture over the entire summer. In a study in Portugal and Spain, LFM of Mediterranean shrub species was also found to be predicted by the Drought Code (DC), a model of soil moisture with a 52-day time lag (Viegas et al. 2001). While we did not find that fog affected LFM loss, it does appear that significant moisture inputs can positively affect LFM. *Adenostoma fasciculatum* maintains relatively high water potentials (Poole and Miller 1975) and continues gas exchange (Redtfeldt and Davis 1996) through most of the summer drought (unpublished data 2^nd^ author). It is still possible that significant summer fog deposition could improve physiological function and increase plant water content.

For *C. megacarpus*, the LFM loss rate was also best predicted by PC IV, which represents soil moisture deficit (DC) and rainfall (Table 3). *Ceanothus* species often have shallow rooting systems (Hellmers et al. 1955) and the model results may reflect *C. megacarpus* responding to changes in moisture in the upper layers of soil as the drought progresses. Yet shallow roots might also allow *C. megacarpus* to access fog water (Figure 5A). This species remains physiologically active during part of the summer drought (at least in some years) (Gill and Mahall 1986) when transpiration should be influenced by VPD. At the CHI field site, LFM loss rate for *C. cuneatus* was best predicted by PC II representing fire weather conditions and soil moisture (Table 3). The Fosberg fire weather index (FFWI) models temperature, relative humidity and wind to measure changes in fire weather conditions (Fosberg 1978). Increased fire weather conditions aggregated over a two-week period (the sampling period of this study) may affect plant physiological function by increasing evapotranspiration and water loss. *C. cuneatus* experiences greater LFM loss under fire weather conditions and responds to lack of water in the top 10 centimeters of soil (modeled in the Drought Code). Although no fog was measured at the CHI field site, *C. cuneatus* is physiologically active during the summer drought (Baker et al. 1982) and may benefit from fog drip when available.

While there is a LFM threshold in the relationship between *A. fasciculatum* LFM and area burned in southern California (Dennison and Moritz 2009), LFM of chaparral species may not be as influential to fire behavior as dead fuels. Additionally, differences in LFM may be more reflective of changes in dry mass and structure instead of water content (Qi et al. 2014). For the chaparral association in the Santa Barbara region, fog was not as frequent during the late summer (Figure 4), possibly explaining why PC III (fog and rain) did not predict LFM loss rate. In California coastal regions near Santa Cruz, fog is more frequent (Lewis et al. 2003) and has been shown to increase soil moisture within a chaparral community (Vasey et al. 2012). With more consistent fog in greater quantities it is possible that fog could affect chaparral LFM loss rates in some years.

Fog effects on LFM were more evident for the sage scrub plant species. This association occurs at lower elevations where fog is more frequent. The LFM loss rates for all three species were best predicted by PC III representing fog deposition and rainfall (Table 3). Additionally, fog alone was a significant predictor of LFM loss for the three species. Fog can reduce evapotranspiration (Fischer et al. 2009), lower growth rates with reduced insolation (Williams et al. 2008, Carbone et al. 2013), and provide a direct source of water for plants (Limm et al. 2009). These effects can positively influence LFM and reduce the dry down rate. The two drought-deciduous species rely heavily on summer water availability to maintain photosynthesis and reduce leaf loss (Poole and Miller 1975). Condensation as dew may also influence water content in sage scrub species as these can be important water resources in arid ecosystems (Agam and Berliner 2006). Though this study was unable to measure dew formation, condensed water vapor may be important for all three sage scrub species.

Summer fog could reduce plant flammability for many species in the sage scrub plant association. This is important as the sage scrub association tends to occur where much of human habitation lies (Davis et al 1994). Humans are the primary cause of ignitions in southern California and are increasing fire frequency in shrubland ecosystems (Syphard et al. 2007). Thus, although ignitions are likely to be high because of human contact with this association, the sage scrub species may experience less successful ignition and lower fire spread rates due to fog influence on live fuel condition.

### Conclusion

This study has shown direct fog uptake is rare, but more importantly, fog deposition reduces the rate of summertime water loss for sage scrub species and lowers their flammability. This influence could become very strong when and if fog inundation is frequent and in high quantity. Although the summer seasons of this study were not particularly foggy compared to fog deposition reported elsewhere in California (Sawaske and Freyburg 2015, Clemesha et al. 2016), fog did reduce LFM loss rates for the sage scrub species. This buffering effect on LFM during the fire season may alter fire behavior across coastal plant associations. By contrast, LFM of higher elevation shrub species that comprise chaparral, were not affected by fog deposition. However, these plant communities did not experience high fog inundation during the late summer, when LFM was lowest. In more northern California coastal shrublands (both sage scrub and chaparral), fog is more consistent and could have an impact on LFM. For many Mediterranean-type climate regions, fog can occur frequently during the dry season, and thus current models of fire danger and LFM in such habitats should include fog deposition and low cloud occurrence.

## Acknowledgements

This work was supported by the UC Natural Reserve system, the California Native Plant Society, UCSB, NSF Doctoral Dissertation Improvement Grant Award # DEB-1311605, the Schuyler Endowment to C. D’Antonio, and an Isotope Inter-University Training for Continental-scale Ecology Fellowship. Water, soil and plant material were collected thanks to El Capitan State Park, Coal Oil Point Reserve, Sedgwick Reserve and Larry and Katy Power’s homestead. Thanks to Keely Roth and Sara Baguskas for extensive knowledge and assistance in conducting fieldwork. Thanks to Dan Okamoto and Mark Wilber for assistance with data analysis. Many thanks to D. Roberts and M. Moritz for their helpful comments on the manuscript.

## Author contributions

N.E. and C.D. developed the research design. N.E. carried out research and analyzed data. C.S. provided consultation and assistance with stable isotope analysis. N.E. wrote and C.D. and C.S. edited the manuscript.

---

**Appendix I.**
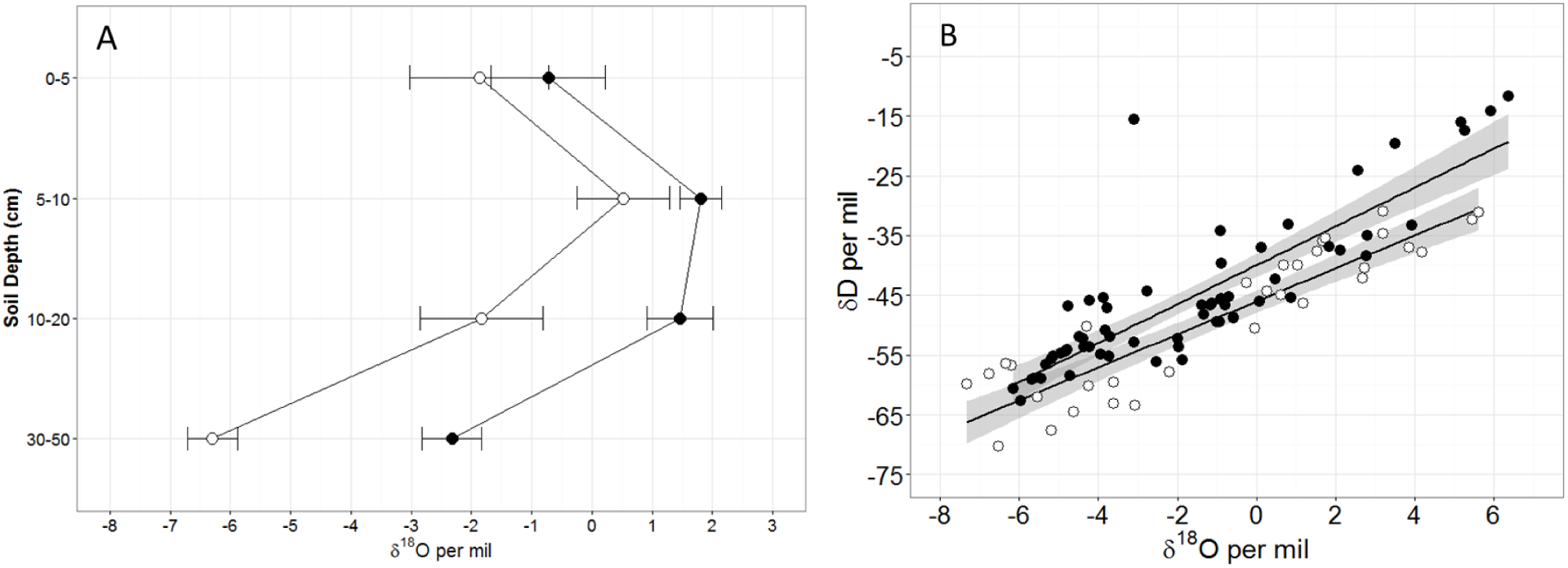
(A) Soil water oxygen isotope ratio values with depth for sage scrub coastal (SSC2, Black) and chaparral coastal (CHC, White) field sites. Points represent the mean value with standard error bars. (B) Soil water evaporation fractionation with linear regressions for each plant association (SSC2 in black, CHC in white). The slope for SSC2 is 3.33 and the slope for CHC is 2.76. The grey area represents the 95% confidence interval around each regression line. The evaporation correction lines were developed using the methods detailed in Corbin et al. (2005). The slopes for the SSC2 and CHC field sites are comparable to the slope of 3.5 found by Corbin et al. (2005), slopes ranging from 2–5 for experimental soil water evaporative experiments using dune soils (Allison 1982) and a slope of 4.0 in a Mexican aquitard (Ortega-Guerrero et al. 1997). These three studies and the slopes found in this study are derived from non-saturated soils and lower than those resulting from saturated soils or open water bodies (Gonfiantini 1986, Barnes and Turner 1998). Variation in the slope is expected and to account for this, plant water samples were corrected using the slope and the 95% confidence interval of each slope before running the MixSIAR mixing model analysis. This resulted in a conservative estimate of proportional fog water in plant tissue for all study species.

**Appendix II.**
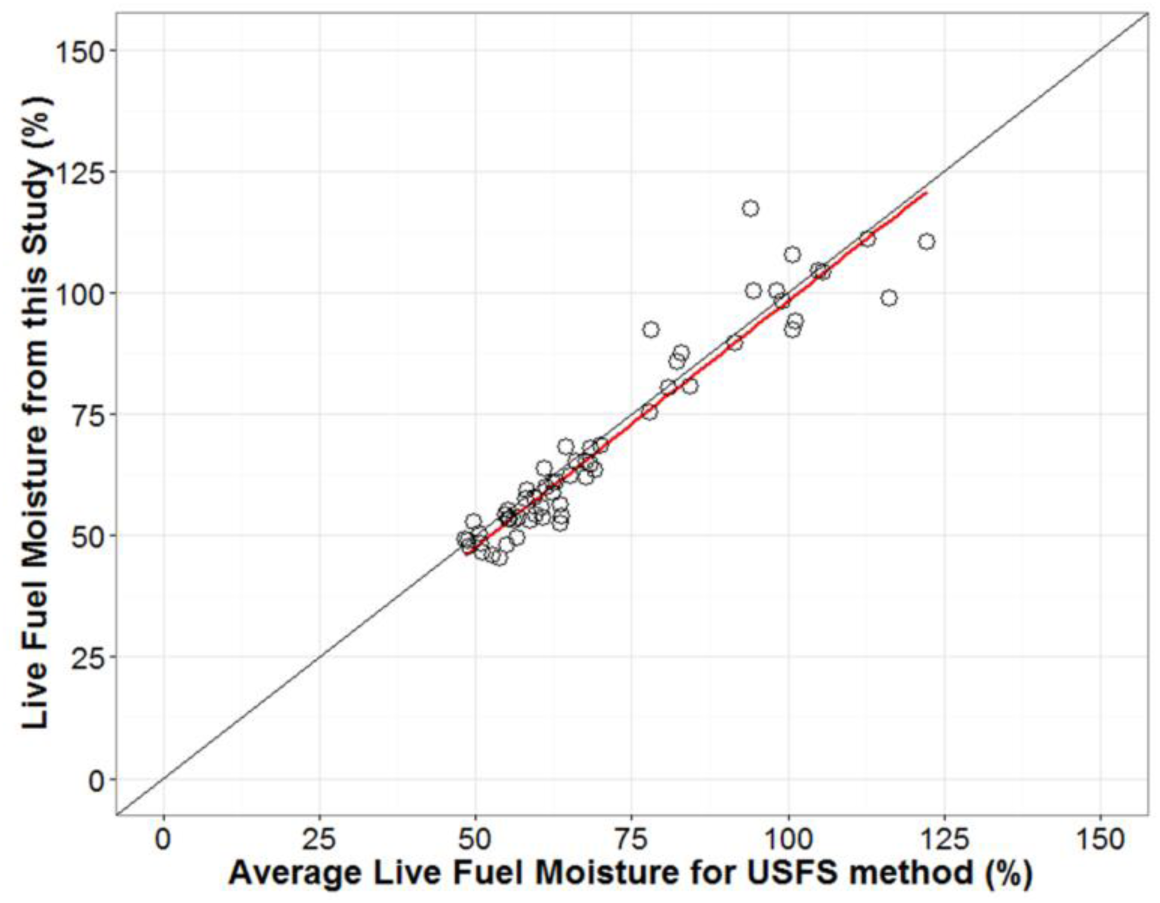
Comparing the LFM method used in this study to the average of new and old LFM material sampled per the USFS method for *A. fasciculatum* (Countryman and Dean 1979). The points represent paired samples from the same shrub individuals. The black line is the 1 to 1 relationship between the two methods and the red line is a linear regression of the two methods (“LFM for this study” = 0.9065 ^*^ “Average LFM using the USFS method” + 0.0848, r^2^ = 0.9184). This figure indicates that in *A. fasciculatum*, the LFM sampling method for this study is very similar to the average of new and old LFM using the USFS method.

**Appendix III.**
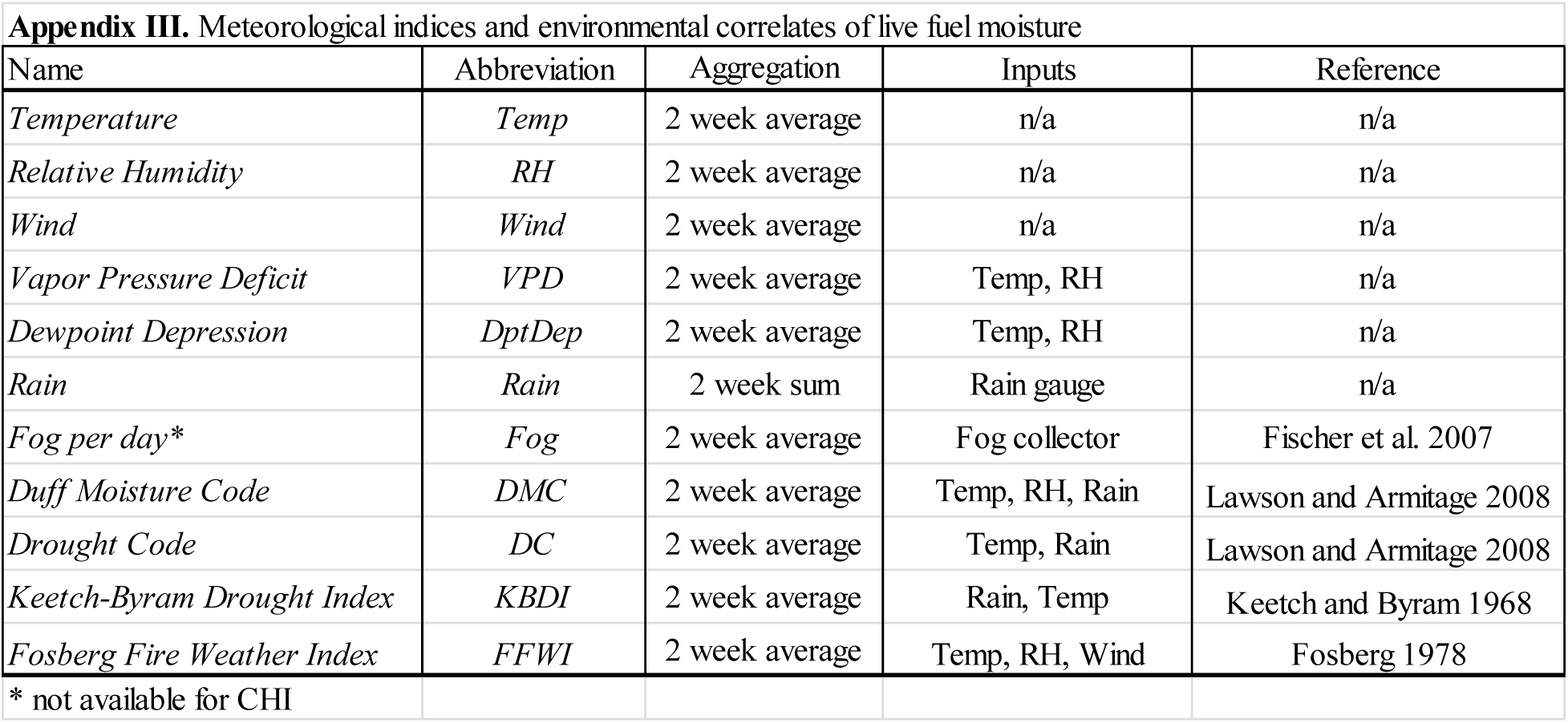
Mean temperature, relative humidity and vapor pressure deficit were used in the linear mixed effects model instead of maximums and minimums because all three were correlated within each metric. The indices and environmental factors were aggregated over two weeks prior to each date of the study. These correlates were decomposed into principal components for the field sites SSC1, SSC2, CHC and SSI. A separate PCA was performed for the CHI field site as no fog deposition data was available.

**Appendix IV.**
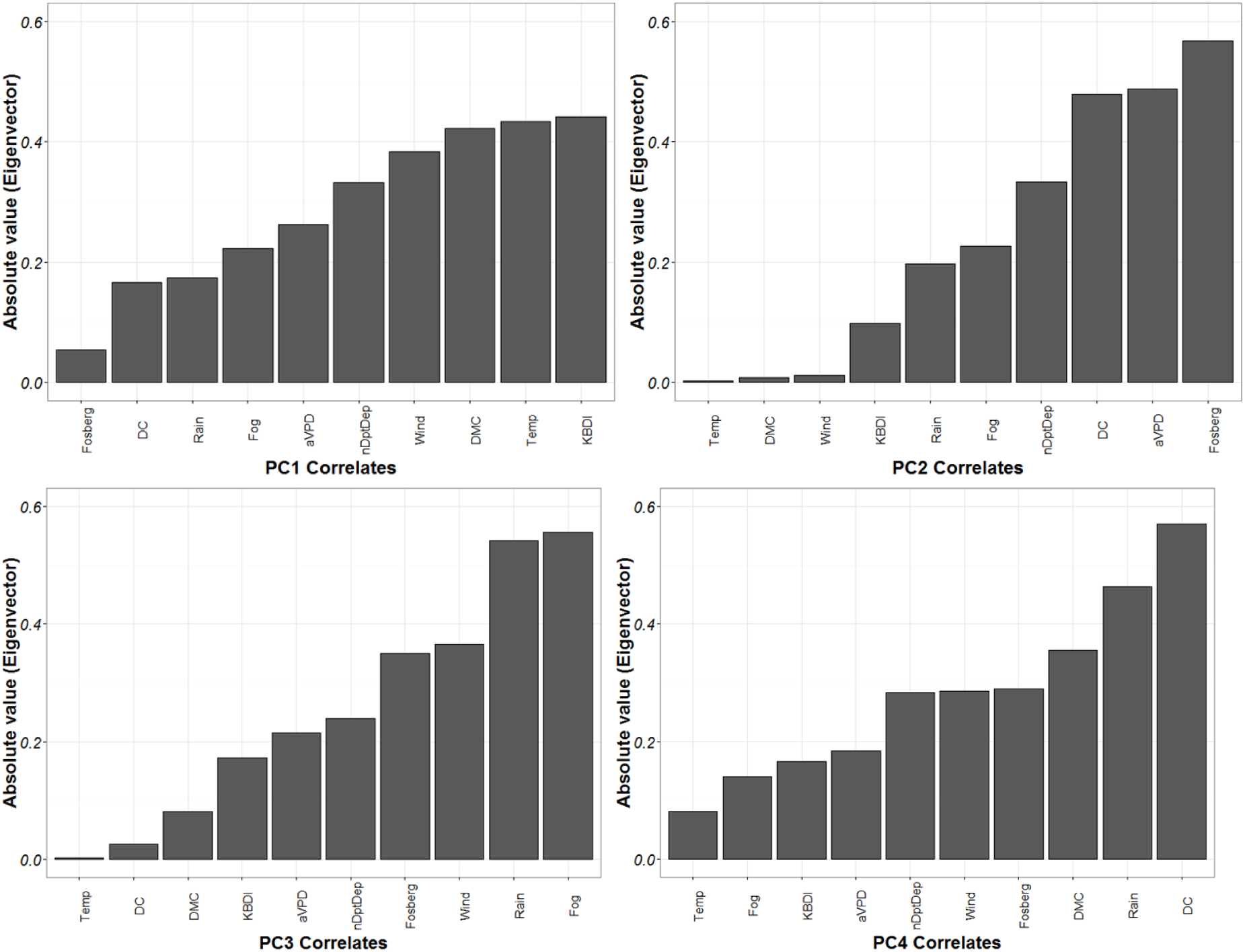
Principal Component loadings for field sites SSC1, SSC2, CHC and SSI. There were four principal components that comprised 87.7% of the variance.

**Appendix V.**
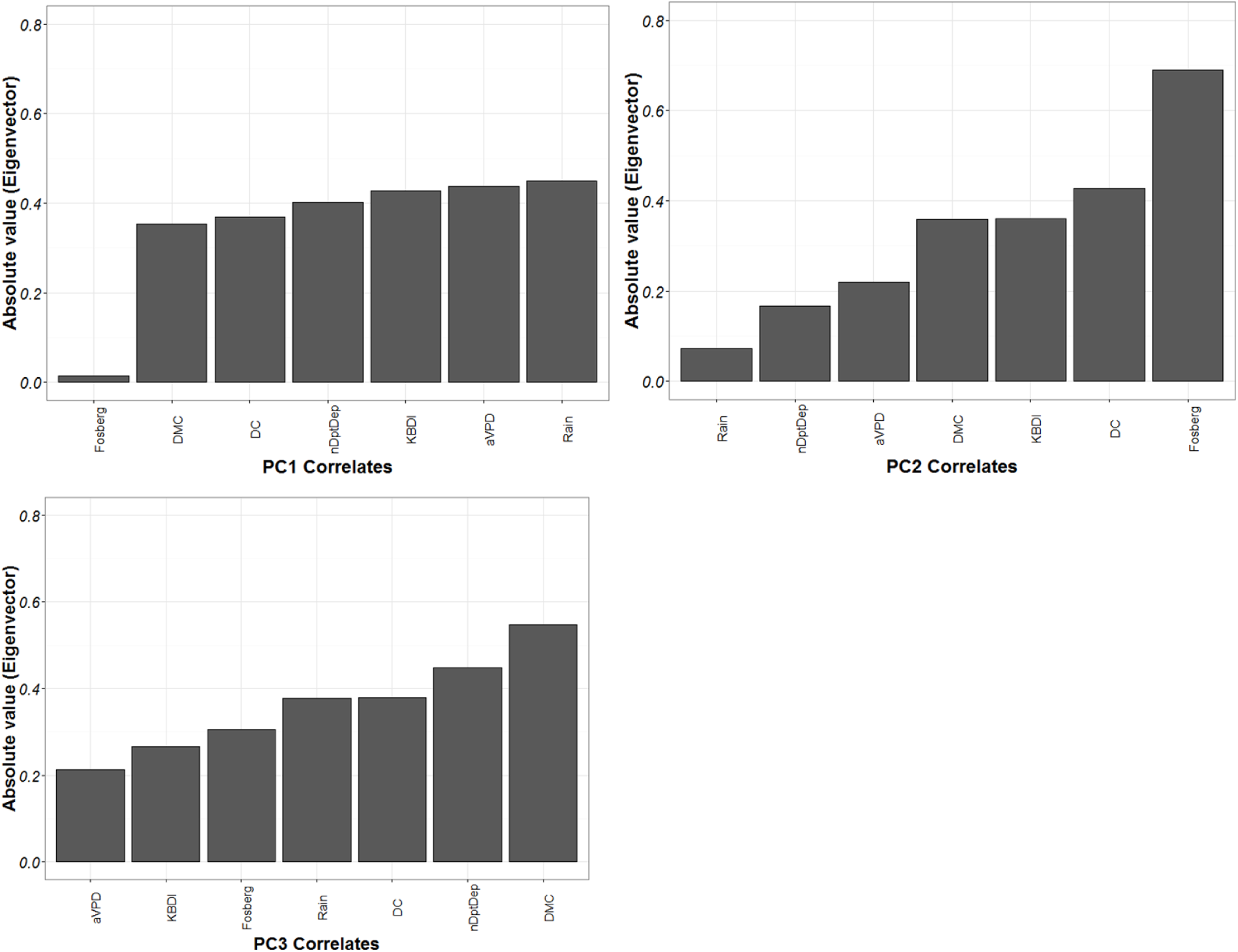
Principal Component loadings for the CHI field site. There were three principal components that comprised 91.3% of the variance.

